# Why is the Omicron main protease of SARS-CoV-2 less stable than its wild-type counterpart? A crystallographic, biophysical, and theoretical study of the free enzyme and its complex with inhibitor 13b-K

**DOI:** 10.1101/2024.03.04.583178

**Authors:** Mohamed Ibrahim, Xinyuanyuan Sun, Vinicius Martins de Oliveira, Ruibin Liu, Joseph Clayton, Haifa El Kilani, Jana Shen, Rolf Hilgenfeld

## Abstract

During the continuing evolution of SARS-CoV-2, the Omicron variant of concern emerged in the second half of 2021 and has been dominant since November that year. Along with its sublineages, it has maintained a prominent role ever since. The Nsp5 main protease (M^pro^) of the Omicron virus is characterized by a single dominant mutation, P132H. Here we determined the X-ray crystal structures of the P132H mutant (or O-M^pro^) as free enzyme and in complex with the M^pro^ inhibitor, the alpha-ketoamide **13b-K**, and we conducted enzymology, biophysical as well as theoretical studies to characterize the O-M^pro^. We found that O-M^pro^ has a similar overall structure and binding with **13b-K**; however, it displays lower enzymatic activity and lower thermal stability compared to the WT-M^pro^ (with “WT” referring to the original Wuhan-1 strain). Intriguingly, the imidazole ring of His132 and the carboxylate plane of Glu240 are in a stacked configuration in the X-ray structures determined here. The empirical folding free energy calculations suggest that the O-M^pro^ dimer is destabilized relative to the WT-M^pro^ due to the less favorable van der Waals interactions and backbone conformation in the individual protomers. The all-atom continuous constant pH molecular dynamics (MD) simulations reveal that His132 and Glu240 display coupled titration. At pH 7, His132 is predominantly neutral and in a stacked configuration with respect to Glu240 which is charged. In order to examine whether the Omicron mutation eases the emergence of further M^pro^ mutations, we also determined crystal structures of the relatively frequent P132H+T169S double mutant but found little evidence for a correlation between the two sites.

## Introduction

In December 2019, the severe acute respiratory syndrome outbreak caused by a novel coronavirus initiated a global epidemic of hitherto unknown dimension. The new coronavirus is ∼82% identical to SARS-CoV at the amino-acid level; hence it is referred to as SARS-CoV-2 [1, 2]. Globally (as of August 2023), there have been over 770 million confirmed cases of SARS-CoV-2 infection and 6.9 million deaths [3]. In November 2021, the lineage BA.1 (earlier B.1.1.529 [4]), the Omicron variant, arose in South Africa and Botswana, and since, new sublineages are springing off from this Omicron variant [5, 6].

In the host cell, the viral proteins are translated from the viral RNA into two huge polyproteins (pp1a, pp1ab), which are processed into 16 nonstructural proteins (Nsps) by the main protease (M^pro^) and the papain-like protease (PL^pro^) [7]. The M^pro^ cleaves the polyproteins at 11 sites that contain the recognition sequence Leu (in one case: Phe)-Gln↓(Ser, Ala, in one case: His) (↓ marks the cleavage site).

M^pro^ is a homodimeric cysteine protease, and each monomer comprises three domains. Domains I and II (residues 10 to 99 and 100 to 182, respectively) consist of β-barrels. A cleft between the two domains harbors a number of subsites (S1’ and S1-S5) that provide the binding site for the substrate. Unlike most proteases of the host, the S1 pocket has a substrate preference for glutamine, which makes M^pro^ an attractive antiviral drug target [8]. Finally, domain III (residues 198 to 303) is a cluster of five α-helices crucial for regulating the dimer formation of the enzyme. Dimerization of the M^pro^ monomer is necessary for the catalytic activity as it allows the interaction between the N-finger (residues 1-9) of one protomer with Glu166 of the second protomer. The N-terminus/Glu166 interaction is essential for forming the S1 subsite [9].

According to GISAID [10], the Omicron variant started replacing the Delta variant around December 2020. By the end of 2021, it became the dominant strain globally, and its evolution is still ongoing, with several main lineages (BA.1, BA.2, BA.4, BA.5, BQ.1, XBB, XBB.1.5, EG.1, EG.5) and their sublineages [10, 11]. The omicron variant accumulated up to 50 mutations throughout the genome [11]. One of the mutations of concern, P132H, is in the Nsp5, affecting the main protease. Such mutation raises concern regarding drug resistance against the available inhibitors for the M^pro^. Structurally, the mutation is located ∼21 Å away from the active site of the M^pro^. However, the P132H mutation site is located at the interface of domains II/III within the same protomer where it participates in forming the so-called distal site, along with Arg131, Thr196, Asp197, Thr198, Asn238, Tyr239, Leu287, and Asp289. Several studies suggest an allosteric effect of this pocket and analyze its druggability with promising scores [12,13]. Further study proposes a long-distance dynamic communication between the distal and active sites that might influence the catalytic activity [14].

One concern is that subtle structural changes accompanying single-site mutations may ease the emergence of secondary mutations that could reduce the affinity of the few approved inhibitors (nirmatrelvir, ensitrelvir) for the M^pro^ and thereby cause resistance towards these drugs. Therefore, we investigate the effect of the mutation, P132H, as well as of the double mutation, P132H+T169S, on the binding of our M^pro^ inhibitor **13b-K** [9, 15] using X-ray crystallography. This secondary mutation is an occasional companion of P132H, and we ask the question whether its emergence is favored by the primary mutation. In addition, we perform molecular dynamics (MD) simulations to understand the possible effects of the P132H mutation on the conformation and structural stability of the M^pro^. We further investigate the residues most influenced by the P132H mutation, near or at the active site, that might be subject to future potential accumulated mutation, as possibly exemplified by T169S. One focus of our study is the protonation state of His132 and its potential hydrogen bonding with Glu240.

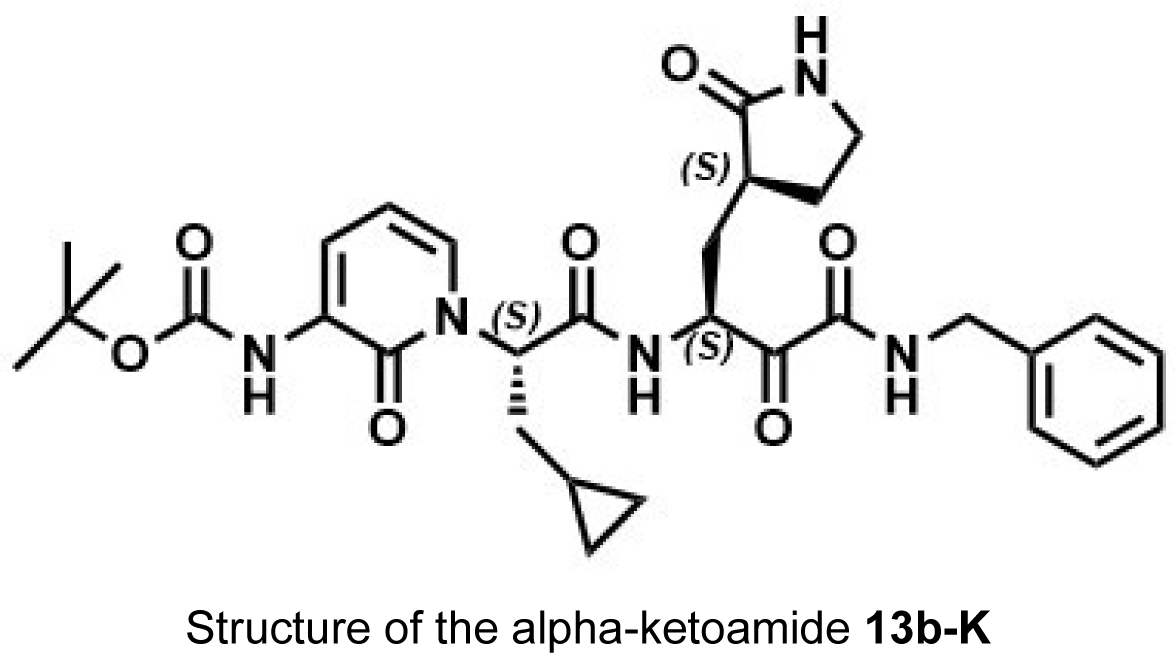

## Results and Discussion

### Enzymatic activity and thermal stability of the Omicron M^pro^

A well-established Förster resonance energy transfer (FRET) assay using a fluorescent substrate, which harbors the cleavage site (indicated by the arrow, ↓) of SARS-CoV-2 M^pro^ (Dabcyl-KTSAVLQ↓SGFRKM-E(Edans)-NH_2_; Biosyntan), was used to evaluate the enzymatic activity of WT-M^pro^ and the two O-M^pro^s (P132H and P132H+T169S). The kinetics of peptide cleavage by the O-M^pro^s is considerably slower compared to that of the wild-type enzyme. We determined the k_cat_/K_m_ for the wild-type enzyme as 6752 ± 1304 M^-1^ s^-1^. In our hands, the k_cat_/K_m_ for the O-M^pro^ is 4915 ± 1278 M^-1^ s^-1^, i.e. 27% lower. This difference is mainly due to k_cat_, which is 0.58 s^-1^ for the WT and 0.40 s^-1^ for the O-M^pro^, indicating that the mutation does not appreciably affect the binding of the substrate but the velocity of its conversion. k_cat_/K_m_ is nearly the same for the P132H+T169S double mutant, but with slightly lower k_cat_ (0.35 s^-1^), compensated by a reduction in K_m_ (72 μM). (Figure 1A, Table 1). These results are largely in agreement with data presented by Greasley et al. [16] and Chen et al. [17], both of whom report a k_cat_/K_m_ reduction by ∼34% for the P132H mutant, but at variance with Sacco et al. (∼9% reduction in k_cat_/K_m_) [18], Lin et al. (∼3% reduction) [19], and Ullrich et al. (∼43% increase) [20].

**Figure 1.**
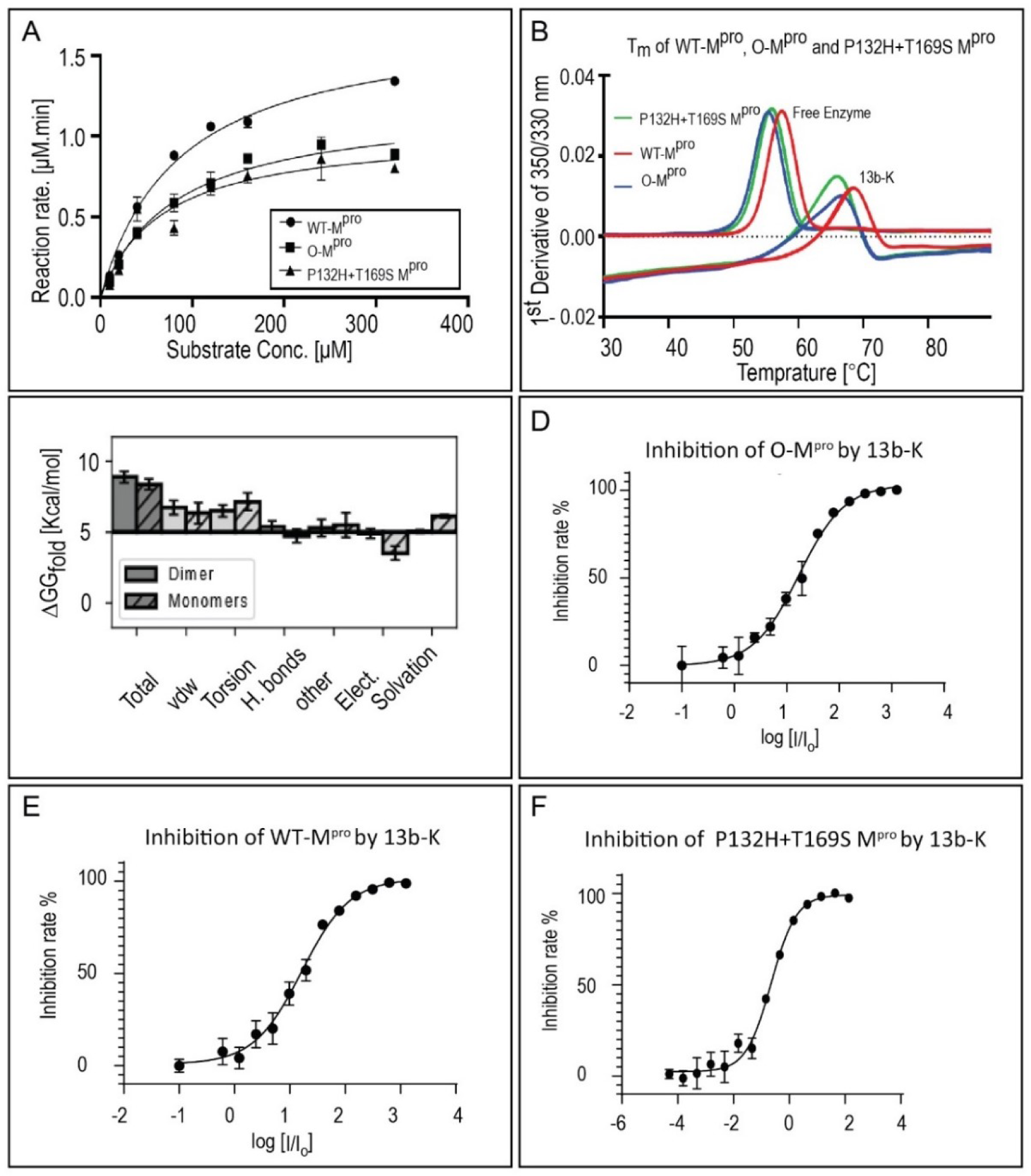
Biochemical and empirical stability characterization of O-M^pro^. (**A**) The Michaelis-Menten kinetics of WT-M^pro^, O-M^pro^, and P132H+T169S double mutant. **(B)** Melting temperature (T_m_) measurements using NanoDSF for WT-M^pro^(red lines), O-M^pro^ (blue lines), and P132H+T169S double mutant (green lines). The measurements were repeated using the same assay to determine the T_m_ after incubation with the **13b-K** inhibitor, with identical color coding. **(C)** Folding free energy difference (ΔΔ*G*_*fold*_) between the WT-M^pro^ and O-M^pro^ dimers (solid), and monomers (striped) calculated from the Rosetta empirical energy functions. Positive ΔΔ*G*_*fold*_ indicates a decreased stability. The total ΔΔ*G*_*fold*_ is broken down into van der Waals (VdW), torsion angle (Torsion), hydrogen bonds (H. bonds), electrostatic interactions (Elect.), solvation free energy (Solvation), and other (Other) terms. Monomer terms are the sums of two monomer energies. **(D, E, F)** The inhibition of O-M^pro^ (D), WT-M^pro^ (E), and P132H+T169S double mutant (F) by **13b-K**.

**Table 1.** Summary of the kinetic parameters and melting temperatures of the M^pro^-WT, O-M^pro^ and P132H+T169S double mutants.

| M<sup>pro</sup> | k<sub>cat</sub><br>(s<sup>-1</sup>) | K<sub>m</sub><br>(μM) | k<sub>cat</sub>/K<sub>m</sub><br>(M<sup>-1</sup> · s<sup>-1</sup>) | Activity<br>(%) | T<sub>m</sub> of<br>M<sup>pro</sup> | T<sub>m</sub> of M<sup>pro</sup><br>with 13b-K |
| --- | --- | --- | --- | --- | --- | --- |
| WT | 0.58 ±<br>0.04 | 85.11 ±<br>14.06 | 6752.05 ±<br>1303.87 | 100.0 | 57.44 ±<br>0.03 | 68.41 ±<br>0.12 |
| O-M<sup>pro</sup> | 0.40 ±<br>0.04 | 81.45 ±<br>19.55 | 4915.08 ±<br>1278.03 | 72.8 | 55.33 ±<br>0.03 | 66.41 ±<br>0.07 |
| P132H+T169S | 0.35 ±<br>0.04 | 71.91 ±<br>24.59 | 4927.46 ±<br>1776.58 | 73.0 | 55.88 ±<br>0.05 | 65.82 ±<br>0.10 |

The inhibitory activities of our α-ketoamide **13b-K** [9, 15] versus SARS-CoV-2 O-M^pro^, the double mutant P132H+T169S, and WT-M^pro^ were determined to assess its activity against these enzyme species. The half-maximal inhibitory concentrations (IC_50_) of **13b-K** against WT-M^pro^ and the two O-M^pro^s were very similar, 0.164 ± 0.030, 0.171 ± 0.034, and 0.209 ± 0.040 μM, respectively (Figure 1D, 1E, and 1F). No substantial differences can be observed in terms of the potency of inhibition by **13b-K**. In line with this observation, nirmatrelvir (PF-07321332) was reported to have a similar inhibition potency for WT-M^pro^ and O-M^pro^ [16].

We also assessed the thermal stability of the O-M^pro^ mutant and that of the P132H+T169S double mutant by nano-Differential Scanning Fluorometry (nano-DSF). In agreement with earlier reports [18], we find that the “melting temperature” of the O-M^pro^ is reduced by 2.1°C, compared to the WT enzyme. The double mutant, P132H+T169S, appears to be slightly more stable than O-M^pro^ (difference: 0.55 °C). Coincubation of each enzyme sample with our inhibitor **13b-K** increases the melting temperature by 10 – 11 °C, but the differences seen between the various forms of the free enzyme (WT, P132H, P132H+T169S) remain the same in the inhibitor complexes (Figure 1B).

### Structural analysis of the O-M^pro^ and double-mutant free enzymes and their complexes with 13b-K at different pH values

The crystal structure of the free O-M^pro^ enzyme (as crystallized at pH 8.2) was determined at 1.91 Å resolution in monoclinic space group *C2* with one M^pro^ chain (half an M^pro^ dimer) in the asymmetric unit. The complex between O-M^pro^ and **13b-K** was crystallized at two different pH values (8.5, 6.5) in order to visualize pH-dependent local differences in the interaction between Ser1* of one O-M^pro^ protomer and Glu166 of the other protomer. (In order to emphasize the intermolecular character of this interaction, Ser1* will be marked by an asterisk in what follows). Crystals grew in triclinic space group *P1* with four O-M^pro^ chains per asymmetric unit and diffracted X-rays to 2.30 Å (pH 8.5) and 2.48 Å (pH 6.5). In addition, the free enzyme of the double mutant P132H+T169S and its complex with our inhibitor **13b-K** was crystallized (at pH 8.0 and 7.5, respectively) in space groups *C2* and *P2_1_*, respectively, with half an M^pro^ dimer and a complete M^pro^ dimer in the asymmetric unit. These crystals diffracted X-rays to 1.8 Å and 1.7 Å, respectively. All structures were refined to good stereochemistry and yielded reasonable statistics (see Table S1). Structure factors and atomic coordinates have been deposited in the Protein Data Bank with accession codes 8R19, 8R26, 8R0V, 8R24, 8R1Q.

The overall structure of the O-M^pro^ enzymes is very similar to the WT-M^pro^ (Figure 2A), albeit small changes can be observed near the mutation site. Close to His132, there is a salt-bridge between Asp289 and Arg131. This salt-bridge has been suggested to be essential for the intramolecular domain II/domain III interface in SARS-CoV M^pro^ [21]. Both WT-M^pro^ and O-M^pro^ feature this salt-bridge, although the presence of His132, which potentially has a charged side-chain due to its close proximity to Glu240 (see below), might influence it directly or indirectly and reduce the protein stability (Figure 2B). In addition, the three-dimensional structure of the free enzyme of the P132H+T169S double mutant is largely unchanged compared to the O-M^pro^. This mutation has almost no structural effect, as the residue is fully solvent-exposed on the surface of the protein. In contrast, in the structure of the complex between the double mutant and our inhibitor **13b-K**, the loop 167–170 has a different conformation in protomer A. In this structure, there is a flip of the Pro168 peptide bond relative to the other structures and the ψ angle of Pro168 is 148°, whereas in the free enzyme, it is –29°, very similar to that in the wild-type M^pro^. The Pro168 conformation is less clear in protomer B due to the poorly resolved electron density of the loop. In summary, however, there are no substantial structural changes in any of the X-ray structures, neither in the P132H mutant nor in the P132H+T169S double mutant.

**Figure 2.**
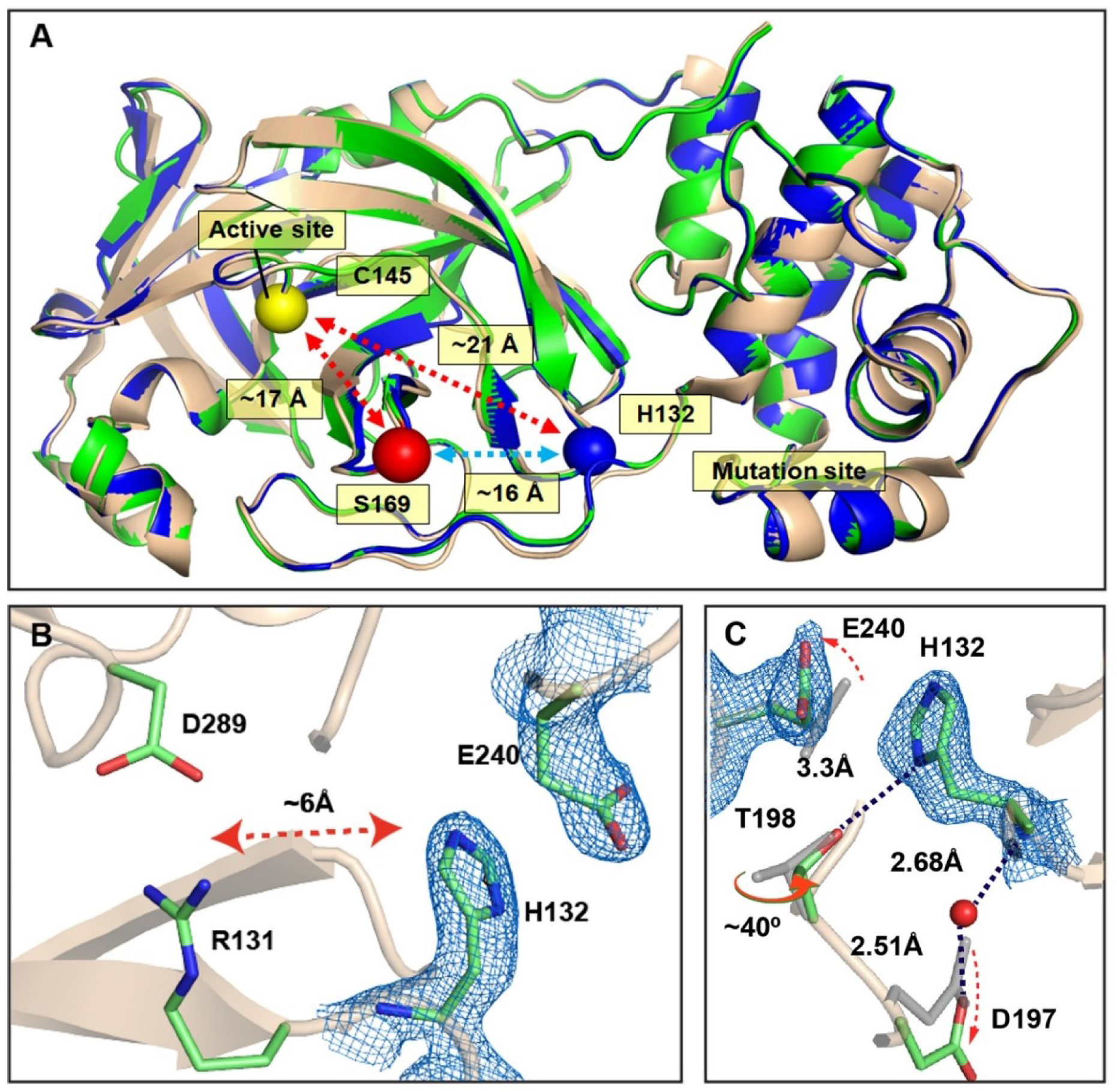
X-ray structure of the free O-M^pro^ and P132H+T169S double mutant in comparison to the free WT-M^pro^. (**A**) Overview of WT-M^pro^ (wheat), O-M^pro^ (blue), and P132H+T169S M^pro^ (green). The active site (yellow ball) is shown in panel A, the mutation sites P132H (blue ball) and T169S (red ball) are also indicated. **(B, C)** A closer view of the O-M^pro^ mutation includes the 2F_o_-F_c_ map contoured at 1.2 σ at His132 (mutation site) and Glu240. (B) shows the close proximity of the mutation site from the Glu289-Arg131 salt-bridge. (C) shows the rotation of Thr198 and the movement of Asp197, with the WT-M^pro^ depicted in gray.

Thr169 is a mutation hot spot of the M^pro^. Its codon is ACU in the Wuhan strain of SARS-CoV-2 but ACG, ACC, or ACA, all coding for Thr, are also found at this site in different M^pro^ sequences (GISAID, [10]). In the T169S mutant of Omicron, the codon is UCU; therefore, the T169S mutation is the result of an A>U transversion. The additional T169S mutation has been found in ∼0.8 % of the O-M^pro^ sequences by mid-April 2023 but seems to experience substantial decline since. Also, there are only very few M^pro^ (<30) sequences that have the T169S mutation but lack the main O-M^pro^ mutation, P132H (all data extracted from GISAID [10]). Based on these data, and on the lack of coupling between His132 and Ser169 (see below), the double mutant P132H+T169S is unlikely to drive further mutation of the Omicron Nsp5 sequence.

Regarding the destabilization of the M^pro^ by the P132H mutation, it has to be considered that His132 occupies a larger space than its ancestor, Pro132, and consequently, the side-chain of Glu240 is shifted to accommodate the newcomer (Figure 2C). In addition, if His132 was charged, it would likely influence the hydrogen-bonding network in the vicinity. Based on this reasoning and the His132-Glu240 distance, we expected to observe an “edge-to-edge” configuration between the two side-chains, in which the imidazole donates a charged hydrogen bond to one of the carboxylate oxygens. Surprisingly, all five X-ray structures, including the free enzyme of both O-M^pro^ and the double mutant, as well as **13b-K**-bound forms obtained at different pH conditions, show a “stacking” configuration between the His132 and Glu240 side-chains, hindering the formation of an H-bond in both protomers. This stacking configuration has also been observed in the X-ray structure of O-M^pro^ in complex with the inhibitor GC376 (PDB accession code 7TOB [18]). On the other hand, the X-ray structure of O-M^pro^ in complex with nirmatrelvir (PDB accession code 7TLL [16]) shows an edge-to-edge configuration, with an imidazole-to-carboxylate distance of 3.2 and 3.8 Å for protomer A and B respectively, indicating the possibility of forming an H-bond. To provide a mechanistic understanding of the unexpected relative orientations of the His132 and Glu240 side-chains and the mutation-induced destabilization, we performed constant-pH and fixed-charge molecular dynamics (MD) simulations as well as empirical protein stability calculations.

### Constant-pH MD simulations reveal that His132 is neutral and prefers the stacking configuration

To understand why His132 and Glu240 engage in a stacking configuration, we performed pH-titration simulations of the free O-M^pro^ using the newly developed GPU-accelerated all-atom continuous constant-pH MD (CpHMD) method [22] in the Amber22 package [23]. The simulations were initiated from the pH 8.5 X-ray structure (which was determined first) and made use of the asynchronous pH replica exchange algorithm for accelerated convergence [24]. Surprisingly, the simulations show that His132 is neutral and Glu240 is charged at pH 7.0, which contradicts our initial hypothesis that His132 is charged and forms a charged H-bond (salt-bridge) with Glu240. Furthermore, the titration of the two residues is coupled. To understand the coupled titration, we calculate the pH-dependent probabilities of the combined protonation states: the doubly protonated state, His132(+)/Glu240(0), the singly protonated states, His132(+)/Glu240(−) and His132(0)/Glu240(0), and the doubly deprotonated state, His132(0)/Glu240(−). Figure 3 demonstrates that at pH 7, the doubly deprotonated state His132(0)/Glu240(−) is predominant for both protomers. Furthermore, as pH decreases, the probabilities of the two singly protonated states increase; however, the His132(0)/Glu240(0) state dominates over the His132(+)/Glu240(−) state (Figure 3A and 3B). Thus, the CpHMD simulations suggest that the singly protonated His132(+)/Glu240(−) pair, which would promote the edge-to-edge configuration through a charged H-bond, is an unlikely state. From the CpHMD simulations, macroscopic pK_a_’s can be obtained by fitting the average number of protons bound to His132 and Glu240 at different pH to a coupled two-proton model [25, 26]. The two stepwise pK_a_’s are estimated as 4.0/6.1 for protomer A and 4.2/5.9 for protomer B (Figure S1). Since the His132(0)/Glu240(0) state is the more probable singly protonated state, the higher pK_a_ can be assigned to Glu240 (6.1 and 5.9 for protomer A and B, respectively) while the lower pK_a_ can be assigned to His132 (4.0 and 4.2 for protomer A and B, respectively). The 0.2-pH unit difference between the pK_a_’s of the two protomers is within the statistical error of the simulations [22].

**Figure 3.**
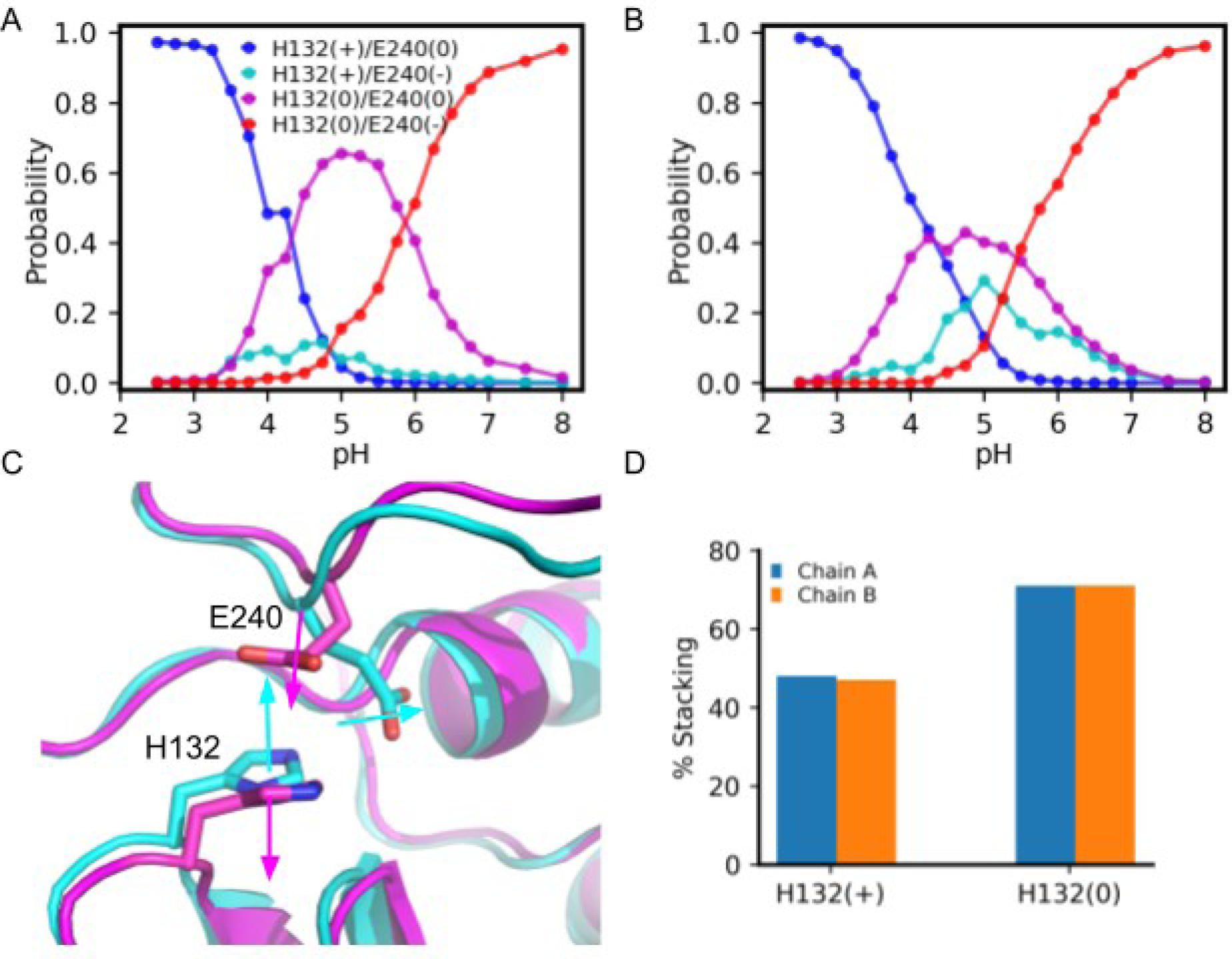
Protonation and configuration states of H132/E240 from the all-atom CpHMD simulations of the free O-M^pro^. (**A, B**) Probabilities of the four protonation states of the H132/E240 pair at different pH in protomer A (A) and B (B). **(C)** Overlay of the MD snapshots of the edge-to-edge (cyan) and stacking (magenta) configuration. His132 and Glu240 side-chains are shown as sticks. The normal vectors of the imidazole ring and the carboxylate plane are shown as arrows. **(D)** Percentage probability of forming the stacking configuration in the presence of charged His132(+) or neutral His132(0) for protomer A (blue) and B (orange).

Next, we examine the effect of the His132 protonation state on the relative His132/Glu240 side-chain conformations using the normal vectors of the imidazole ring of His132 and the carboxylate plane of Glu240 (Figure 3C). The configuration is considered edge-to-edge, if the angle between the two normal vectors is between 60° and 120°, i.e., the imidazole and carboxylate planes are approximately perpendicular, and otherwise stacking (Figure 3C). Based on this definition, the CpHMD simulations demonstrate that His132/Glu240 can adopt either the edge-to-edge or the stacking configuration. The stacking configuration is observed in the X-ray structures obtained in this work and the crystal structure of the O-M^pro^ in complex with GC376 (PDB accession code 7TOB, [18]). On the other hand, the edge-to-edge configuration is found in the X-ray structure of the O-M^pro^ in complex with nirmatrelvir (PDB accession code 7TLL) [16]. Interestingly, the CpHMD data show that the relative His132/Glu240 configuration is dependent on the protonation state of His132. In the presence of His132(+), the probability of forming the stacking configuration is about 50%, i.e., stacking and edge-to-edge configuration are equally probable, whereas in the presence of neutral His132(0), the probability of the stacking configuration is about 70%, i.e., stacking is preferred (Figure 3D). However, since the His132(+) state has a low probability across the simulation pH range (2.5--8), the stacking configuration is always preferred. This is in agreement with the stacking configuration being observed in both of our X-ray structures of O-M^pro^ in complex with **13b-K** at pH 6.5 and 8.5.

### Fixed-charge MD simulations support the preference for the stacking configuration of His132 and Glu240

Although the CpHMD data demonstrate the preference for the stacking configuration, the simulation length was limited. Thus, to confirm the finding, we conducted two runs of 500-ns fixed-charge MD simulations starting from the pH 8.5 X-ray structure of the free O-M^pro^ (PDB accession code 8R19) with His132 in the neutral state. Additionally, four runs of 1-μs fixed-charge MD simulations were performed based on the published X-ray structure with the PDB accession code 7TOB [18] (inhibitor removed), whereby His132 was either charged (two runs) or neutral (two runs). The simulations based on our X-ray structure model show that the imidazole ring of His132 and the carboxylate plane of Glu240 are predominantly stacked, as the angle between the two normal vectors (defined above) is either 30° or 150° (Figure 4A). In contrast, the edge-to-edge state is infrequently sampled, as the angle distribution plot displays a valley between 60° and 120°. Note, the second trajectory gave very similar results (Figure S2). The simulations based on 7TOB [18] confirm that the stacking configuration is favored when His132 is neutral whereas the edge-to-edge configuration is favored when His132 is charged (Figure S3).

**Figure 4.**
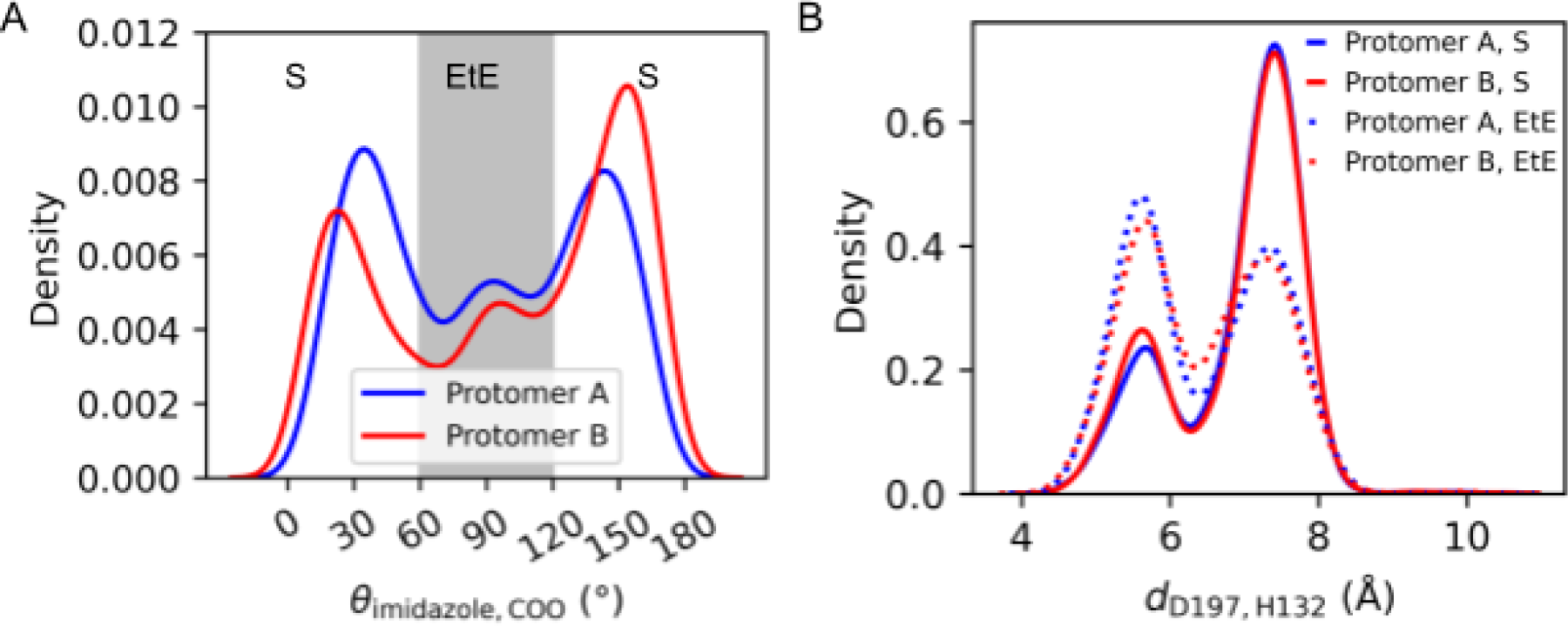
Configuration of H132/E240 and the effect on the water-mediated D197-H132 interaction from the fixed-charge MD simulations. (**A**) Distributions of the angle between the normal vectors of the His132 imidazole and Glu240 carboxylate planes for protomer A (blue) and B (red). The grey area represents the edge-to-edge (EtE) configuration, while the other parts represent the stacking (S) configuration. **(B)** Distributions of the minimum distance between the carboxylate oxygens of Asp197 and the backbone carbonyl oxygen of His132 for the His132/Glu240 stacking (solid lines) and edge-to-edge (dotted lines) configuration. Data for protomer A and B are shown in blue and red, respectively.

### MD analysis suggests that the local electrostatic and H-bond environment of His132 is largely unperturbed

Our X-ray structure of the free O-M^pro^ shows a salt-bridge between Asp289 and Arg131, which is also present in the WT-M^pro^ and has been hypothesized to be important for the intramolecular domain II/III interface in SARS-CoV(−1) M^pro^ [21]. We calculated the probability distribution of the minimum Asp289-Arg131 charge center distance based on the fixed-charged MD, which confirms that the salt-bridge is stable regardless of the His132/Glu240 configuration (the most probable distance is 2.8 Å for both stacking and edge-to-edge configurations, Figure S2 and Figure S4). This is consistent with the CpHMD finding that His132 is neutral, which does not perturb this salt-bridge interaction. Another residue in the vicinity of His132 is Thr198. In our X-ray structure of the free O-M^pro^, the distance from the His132 imidazole nitrogen to the Thr198 hydroxyl oxygen is 3.3 Å (Figure 2C), suggesting that they might form a dynamical H-bond in solution. To examine this possibility in light of the two different His132/Glu240 configurations, we calculated the distribution of the minimum distance from the imidazole nitrogen to the hydroxyl oxygen for the stacking and edge-to-edge configurations of the free O-M^pro^ (Figure S2 and Figure S4). Although there is a minor peak at about 3.0 Å, the major peak (the most probable distance) is at about 5.5 Å for the His132/Glu240 stacking configuration. For the edge-to-edge configurations, the major peak is at about 4.5 Å. Thus, our simulations do not support an H-bond between Thr198 and His132 in O-M^pro^.

### MD analysis suggests that the His132/Glu240 stacking configuration gives space for water to mediate the sidechain-to-backbone interaction between Asp197 and His132

In our crystal structure of the free O-M^pro^, the Oδ1 atom of Asp197 shifts by 2.2 Å and a water molecule appears to mediate an H-bond between the main-chain nitrogen of His132 and the carboxylate of Asp197 (Figure 2C). This water is absent in the X-ray structures of the WT-M^pro^. To examine if this water-mediated interaction is possibly related to the interaction between His132 and Glu240, we calculate the minimum distance between the carboxylate oxygens of Asp197 and the backbone carbonyl oxygen of His132 for the His132/Glu240 stacking and edge-to-edge configurations based on the fixed-charge MD simulations. For both configurations, the Asp197-His132 distance has two most probable values, about 5.5 Å and 7.5 Å (two peaks in the distributions of Figure 4B); however, the relative probability of the two distances is dependent on the His132/Glu240 configuration. When His132 is in the stacking configuration with respect to Glu240, the larger distance (7.5 Å) is sampled with a much higher probability than the smaller distance (5.5 Å); in contrast, when His132 is in the edge-to-edge configuration, the two distances are sampled with equal probabilities (Figure 4B). Since a larger distance between the Asp197 side-chain and the His132 backbone makes space for the entrance of water, the simulation data suggests that the stacking configuration may promote the water-mediated interaction between Asp197 and His132.

### Empirical folding free-energy calculations predict that the P132H mutation destabilizes M^pro^

To understand the mechanism of the stability decrease of O-M^pro^ relative to the WT, we estimate the folding free energy change ΔΔ*G*_*fold*_ for both the dimers and monomers using the ddG_monomer application in the Rosetta software suite [27]. The calculation gives a positive ΔΔ*G*_*fold*_ of 7.8 ± 0.8 kcal/mol, indicating that the P132H mutation destabilizes the M^pro^ dimer, which is consistent with the melting temperature reduction of 2.1°C. The destabilization is mainly driven by unfavorable van der Waals interactions and backbone torsion energies, indicating that the P132H mutation induces steric repulsion and unfavorable conformations (Figure 1C). To test if the dimer interface is involved in the stability change, we examine ΔΔ*G*_*fold*_ of the monomers. If the sum of the monomer ΔΔ*G*_*fold*_ and the individual contributions are the same as the corresponding ΔΔ*G*_*fold*_ of the dimer, the stability changes of the monomers are solely responsible for the stability change of the dimer, i.e., no change in the dimer interface plays a role. On the other hand, if ΔΔ*G*_*fold*_ of the dimer is greater (more positive) than that of the monomers, the difference can be attributed to the destabilization of the dimer interface. Following this reasoning, we examine the differences between the dimer and monomer calculations. The sum of monomer ΔΔ*G*_*fold*_ is 6.8 ± 0.8 kcal/mol; the difference from the dimer ΔΔ*G*_*fold*_is 1.0 kcal/mol, which is within the error bar of 1.1 kcal/mol (Figure 1C). The greatest difference between the individual energy terms of the monomers and the dimer is in the electrostatic and solvation energies. The P132H mutation is predicted to have stabilizing electrostatic (−3.0 ± 1.1 kcal/mol) and destabilizing solvation energies (4.3 ± 1.7 kcal/mol) for the monomers, whereas these terms are nearly unchanged for the dimer. It is noteworthy that the stabilization of electrostatics and destabilization of solvation largely cancel out, making the net difference between the monomers and the dimer negligible. Taken together, these data suggest that the destabilization of O-M^pro^ relative to the WT is mainly due to the destabilization of the individual protomers, and although the electrostatics and solvation of the dimer interface are affected by the mutation, the net effect on the stability of the dimer is negligible.

### The P132H mutation has no negative impact on 13b-K binding

The active site of the M^pro^ is ∼21 Å away from the discussed mutation (Figure 2A). No substantial changes were observed between the overall structures of WT-M^pro^ and O-M^pro^. The RMSD values of the models were calculated for the complete chain against the WT-M^pro^ (PDB 6Y2E [9]), additionally for each domain separately. With four copies (two dimers) of the O-M^pro^–**13b-K** complex in triclinic space group *P1*, such calculations potentially offer interesting results regarding subtle differences between individual M^pro^ complex molecules (such information is not available from the common M^pro^ complexes crystallized in space group *C2*, which has a two-fold axis of symmetry). Thus, a difference in the RMSD of each chain within the model of the O-M^pro^/**13b-K** complex at pH 8.5 is observed. Chains A and C have higher RMSD values (0.43 Å and 0.47 Å) than chain B and chain D (0.37 Å and 0.40 Å; values are for all non-hydrogen atoms). On the other hand, within the model of the O-M^pro^/**13b-K** complex at pH 6.5, the four chains have almost identical RMSD values (0.37 Å, 0.38 Å, 0.37 Å, and 0.37 Å) for chains A, B, C, and D, respectively. For the high-pH form, calculation of RMSD values for the individual amino-acid residues (using the WT-M^pro^ (6Y2E) as a reference) for each of the four chains, A – D, reveals that in specific regions, i.e., in the vicinity of Ser1*, Asn27, Gln74, Pro168, or Thr279, a considerable increase in the RMSD values exists for the A/C chains but not the B/D chains (Figure S6). One of these differences is due to a different arrangement of the interaction between Ser1* and Glu166 in chains A and C, compared to B and D. In the former, the N-terminus (Ser1*) and Glu166 are connected via a salt-bridge, whereas in the latter, this salt-bridge is absent and there is instead an H-bond between the side-chain hydroxyl of Ser1* and the carboxylate of Glu166 (Figures 5E and 5F). The reason for this feature could be that at the pH of crystallization (8.5), the free amino group of Ser1* may be partly deprotonated, as the solution pK_a_ of the N-terminal amino group of a protein is ∼8.0 – 8.5. This may vary in the M^pro^ dimer, as the N-terminal Ser1* is only partially solvent-accessible due to its engagement with Glu166, and further modulation may be caused by the nearby His172 [28]. When the N-terminal Ser1* is deprotonated, the salt-bridge with Glu166 cannot be formed and is replaced by the H-bond between Ser1* Oγ and the Glu166 carboxylate. Along with these changes in the H-bonding pattern, the side-chain torsion angles χ1 and χ2 of Glu166 are in the antiperiplanar *(ap)* / *ap* range in protomer A and in the synclinal *(sc)* / *ap* range in protomer B. This conformational change of the Glu166 side-chain in protomer B enables it to form a new hydrogen bond with His172 Nη2. At elevated pH, each O-M^pro^ dimer would thus consist of one protomer (A or C) comprising the salt-bridge between Ser1* and Glu166 (with Glu166 in *ap/ap* conformation) and one protomer (B or D) lacking the salt bridge (and Glu166 in *sc/ap* conformation). This interpretation is supported by the fact that the crystals of the O-M^pro^/**13b-K** complex grown at pH 6.5 show negligible RMS differences between all four chains, and the salt-bridge is fully formed in all four copies of the Ser1*–Glu166 pair. In the **13b-K** complex of the double mutant, P132H+T169S, which was crystallized at pH 7.5, the situation is comparable to the high-pH form of the P132H single-mutant/**13b-K** complex, i.e. the dimer also comprises one protomer with a salt-bridge between Ser1* and Glu166 and one having the Ser1* Oγ–Glu166 H-bond instead (Figures 5G and 5H).

**Figure 5.**
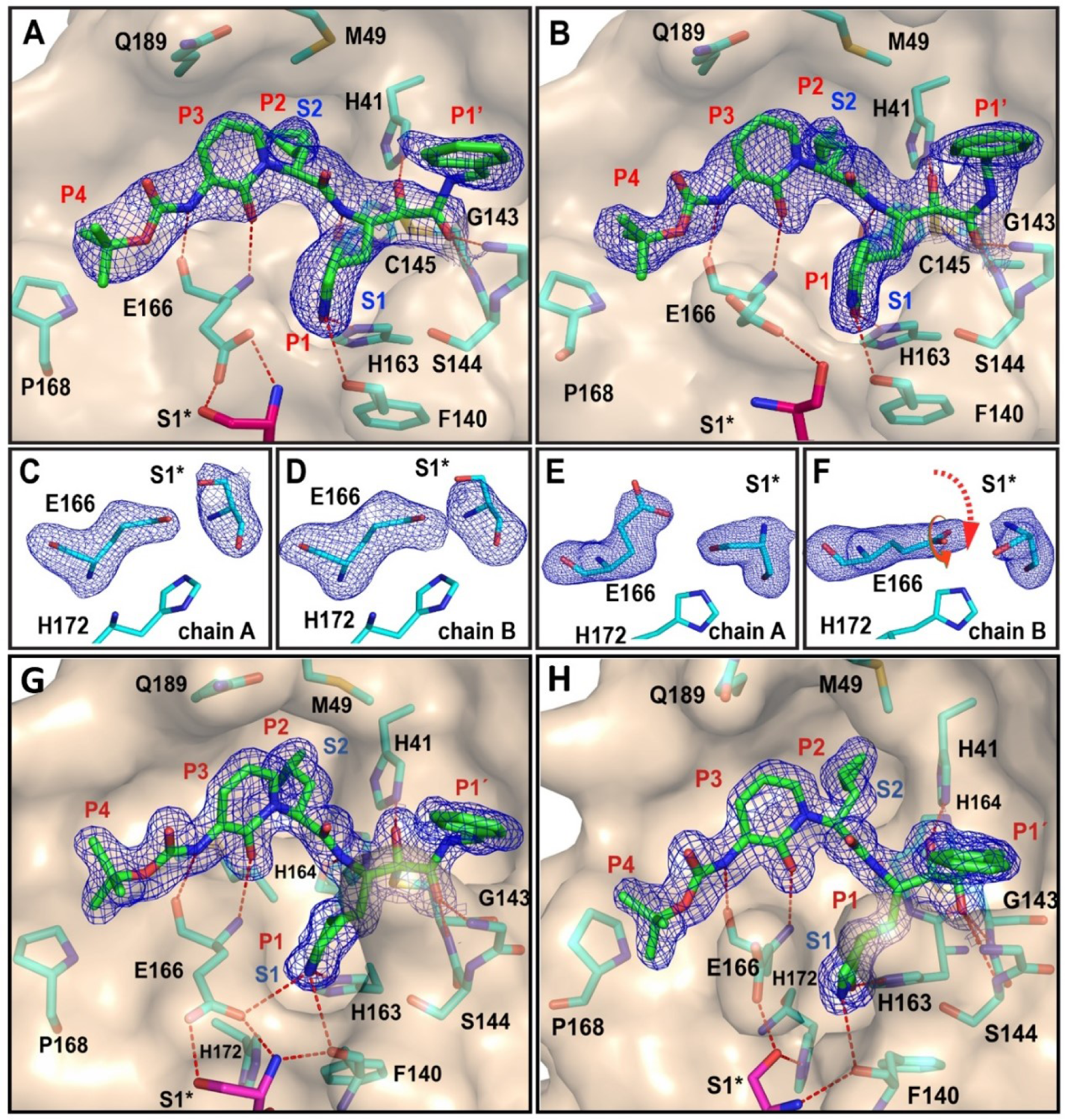
X-ray structures of O-M^pro^/P132H+T169S M^pro^ in complex with 13b-K. (**A, B**) The substrate-binding cleft of the O-M^pro^ crystallized in a complex with **13b-K** at two different pH values, 6.5 (A) and 8.5 (B). The carbon atoms of the inhibitor are colored green, and those of the protein are colored cyan, except for Ser1*, which are in magenta. The binding pockets (S1’-S2) of the protein and the binding moieties (P1’-P4) of the inhibitor are labeled. The 2F_o_-F_c_ map carved around the inhibitor is depicted as mesh and contoured at 1.0 σ. **(C-F)** Interactions of the N-terminus in the two protomers of the same dimer of the pH 6.5 crystal (C and D) and of the pH 8.5 crystal (E and F). Red arrows indicate the difference between the two protomers in the latter case. The represented maps are polder maps calculated in *p*ℎ*enix*. *polde*r [31], by omitting the complete residues of interest, Glu166 and Ser1*, contoured at 3.0 σ. **(G, H)** The substrate-binding cleft of the P132H+T169S double mutant co-crystal structure with **13b-K**, including protomer A (G) and protomer B (H).

The discussed interaction between Ser1* and Glu166 (a salt-bridge at pH 6.5 and either a salt-bridge or an H-bond at pH 8.5) is crucial for forming the S1 pocket (Figure 5A, 5B, 5E, and 5F). We have previously observed similar differences at Ser1* in the crystal structure of the high-pH form of the SARS-CoV-2 WT M^pro^ (PDB 6Y2G, molecule A) [9], although it has to be mentioned here that the originally deposited high-pH structure of the SARS-CoV-2 WT M^pro^ structure had to be corrected, as the interaction between Ser1* and Glu166 had been modelled incorrectly. We have recently applied this correction and uploaded the revised coordinates to the PDB. A similar asymmetry in recognition of the Ser1*–Glu166 pair has also been observed in the neutron diffraction study of the telaprevir–M^pro^ complex described by Kovalesky’s group [29].

On the other hand, the O-M^pro^ model at pH 6.5 behaves differently; all monomers have similar salt-bridge interactions between Ser1* and Glu166 (Figures 5C and 5D). In spite of the discussed differences, it is essential to emphasize that independently of the pH of crystallization, our inhibitor, **13b-K**, binds efficiently to all four protomers without significant structural differences (Figures 5A and 5B). And yet, the structural differences around the Ser1*-Glu166 pair become evident only in the structures of the **13b-K** complexes at elevated pH (≥7.5), not in the free enzyme. This is in agreement with the observation by Kneller et al. [29], who showed by neutron crystallography that binding of the HCV NS3/4A inhibitor telaprevir to the M^pro^ leads to a shift in the protonation status of several histidine residues near the active site, along with the aforementioned conformational change of the side-chain of Glu166 in one of the protomers within the dimer. This phenomenon could well be related to the observed half-site reactivity of the SARS-CoV M^pro^ described by Chen et al. [30].

## Conclusions

In this work, X-ray crystallography, biochemical experiments, and MD simulations are employed to characterize the structure and dynamics of the O-M^pro^ P132H in complex with the α-ketoamide-based covalent inhibitor **13b-K**. The FRET assay for the O-M^pro^ shows that its enzymatic activity is 27% lower than that of the WT-M^pro^. Our kinetic results are in agreement with some reports [16, 17] but at variance from others that describe nearly-equal kinetics for O-M^pro^ and WT-M^pro^ [18–20]. The nano-DSF results show that the O-M^pro^ has substantially lower thermal stability than the WT-M^pro^, in agreement with previously reported characterizations of the O-M^pro^ [17, 18]. His132 is located at the intramolecular domain II/III interface and in the close proximity of the charged residue Glu240. In addition, it is close to the Asp289-Arg131 salt-bridge. Thus, we originally hypothesized that the P132H mutation would introduce a new salt-bridge with Glu240 and perturb the local electrostatic and H-bonding networks. However, the crystallographic results at pH 6.5 and 8.5 show that the structures of O-M^pro^ and WT-M^pro^ are very similar, with the Asp289-Arg131 salt-bridge remaining intact. Curiously, His132 and Glu240 are in a stacking arrangement between the imidazole ring and carboxylate plane as opposed to the edge-to-edge configuration if His132 was charged and donated an H-bond to the carboxylate of Glu240. The constant-pH MD simulations find that the titrations of His132 and Glu240 are coupled and His132(0)/Glu240(−) is the most probable protonation state, which explains why the stacking configuration is observed in the X-ray structures. The fixed-charged MD simulations confirm that the stacking configuration is favored over the edge-to-edge configuration for the His132(0)/Glu240(−) pair. Moreover, the stacking configuration results in more space for water to mediate an H-bond between the backbone of His132 and the side-chain of Asp197, which is consistent with the X-ray structures. To explain the observed thermal stability difference between the O-M^pro^ and WT-M^pro^, empirical folding free-energy calculations are employed, which show that the O-M^pro^ is less stable due to unfavorable van der Waals and backbone torsion energies, suggesting that the P132H mutation induces steric repulsion and unfavorable conformations.

We do not observe any negative impact of the P132H mutation on the binding of the inhibitor **13b-K**. In addition, the IC_50_ measurements show similar values compared to the WT-M^pro^. In all structures, **13b-K** shows covalent binding to Cys145 and a clear electron density in all binding sites. The assessment of the RMSD against WT-M^pro^ (6Y2E) [9] shows that at pH 8.5, each dimer of the two in the asymmetric unit contains one protomer that has a higher RMSD value than its partner. This difference is manifested mainly in the interaction of the N-terminus, Ser1*, with Glu166. A similar phenomenon has been seen earlier in our high-pH crystal structure of the **13b-K** complex of WT-M^pro^ (PDB entry 6Y2G) [9] and in the neutron structure of the WT-M^pro^ in complex with telaprevir [29].

Interestingly, the structure of the **13b-K** co-crystal at pH 6.5 does not show similar behavior. We also determined the crystal structure of the double mutant P132H+T169S. T169S seems to be the most frequent companion of the P132H mutation (occurring in ∼0.8 % of the O-M^pro^ sequences, although it appears to experience some decline), but we did not find indications for a coupling between the two mutations, suggesting that they arose independently of each other. Overall, further investigation is needed to understand better the potential danger from Omicron-based M^pro^ mutations that might arise and lead to drug resistance.

## Materials and Methods

### Cloning, gene expression, and purification of SARS-CoV-2 M^pro^ and mutants

The P132H mutation was inserted by overlap extension-PCR reaction. A pair of special primers, P132H_forward (AGTGTGCTATGCGTCATAACTTCACGATCAAA; the underlined sequence corresponds to the mutated P132H codon) and P132H_reverse (TTTGATCGTGAAGTTATGACGCATAGCACACT) was designed.

The first PCR reaction was performed to generate two splice fragments containing a 5′ overhang. The wild-type M^pro^ construct was used as a template. The second PCR joined these two spliced fragments to generate the PCR product encoding the P132H-mutated M^pro^. The amplified PCR product was digested with *BamHI* and *XhoI* and ligated into the vector PGEx-6p-1 (GE Healthcare) digested with the same restriction enzymes. The mutation T169S was inserted using another pair of primers and the M^pro^ P132H construct was used as template. The T169S primers were, T169S-forward: ACATGGAATTGCCG<u>AGC</u>GGTGTACAT; T169S-reverse: ATGTACACC<u>GCT</u>CGGCAATTCCATGT.

The gene sequence of the mutated M^pro^ constructs were verified by sequencing (MWG Eurofins). The expression and purification of sequence-verified SARS-CoV-2 M^pro^ wild-type and mutated M^pro^ were performed as described previously [9]. The proteins were concentrated to around 15 mg/ml in crystallization buffer (20 mM Tris, 150 mM NaCl, 1 mM EDTA, 1 mM DTT, pH = 7.5).

### Determination of protein stability by nano-differential scanning fluorimetry (nanoDSF)

Thermal-shift assays of SARS-CoV-2 M^pro^ and its mutants, or the protein-inhibitor complexes, were carried out using the nanoDSF method as implemented in the Prometheus NT.48 (NanoTemper Technologies). The nanoDSF method is based on the autofluorescence of tryptophan (and tyrosine) residues to monitor protein unfolding. As the temperature increases, the protein will unfold and the hydrophobic residues get exposed; the ratio of autofluorescence at wavelengths 350 nm and 330 nm will change. The first derivative of 350/330 nm can be used to determine the melting temperature (T_m_). The WT or mutant proteins at 10 µM and **13b-K** at 100 µM were mixed in a running buffer (20 mM HEPES, 120 mM NaCl, 0.4 mM EDTA, 4 mM DTT, 20% glycerol, pH 7.0), in a final volume of 15 µL for each measurement. 10 µM of proteins were diluted in the same buffer without inhibitor, to prepare the free enzyme measurement. The free enzyme samples were supplemented with 1% DMSO to achieve the same final concentration as in the protein-inhibitor mixtures. Then all samples were loaded into Prometheus NT.48 nanoDSF Grade Standard Capillaries (PR-C002, NanoTemper Technologies), the fluorescence signal was recorded under a temperature gradient ranging from 25 to 90^◦^C (flow rate of 0.5^◦^C·min^−1^). The melting curve was drawn using GraphPad Prism 7.0 software; the values of the first derivative of 350/330 nm were displayed on the Y axis. The melting temperature (T_m_) was calculated as the mid-point temperature of the melting curve using the PR.ThermControl software (NanoTemper Technologies).

### Enzyme kinetics assays

The enzymatic activity and IC_50_ of SARS-CoV-2 M^pro^ or mutants was measured using a FRET assay [9]. A fluorescent substrate harboring the P1/P1’ cleavage site (indicated by ↓) of SARS-CoV-2 M^pro^ (Dabcyl-KTSAVLQ↓SGFRKM-E(Edans)-NH_2_; Biosyntan) and buffer composed of 20 mM HEPES, 120 mM NaCl, 0.4 mM EDTA, 4 mM DTT, 20% glycerol, pH 7.0 was used for these assays. The fluorescence signal was monitored at an emission wavelength of 460 nm with excitation at 360 nm, using a SPARK Multimode Microplate Reader (TECAN).

To measure the enzymatic activity, 10 µL of SARS-CoV-2 M^pro^ WT or mutant at a final concentration of 50 nM was initially pipetted into a 96-well plate containing pre-pipetted 60 µL of reaction buffer. Subsequently, the reaction was initiated by addition of 30 µL of the substrate dissolved in the reaction buffer to 100 µL final volume, at different final concentrations varied from 10 to 320 µM (10, 20, 40, 80, 120, 160, 240, 320 µM). A calibration curve was generated by measurement of varied concentrations (from 0.04 to 6250 nM) of free Edans, with gain 80 in a final volume of 100 µL reaction buffer. Initial velocities were determined from the linear section of the curve, and the corresponding relative fluorescence units per unit of time (ΔRFU/s) was converted to the amount of the cleaved substrate per unit of time (µM/s) by fitting to the calibration curve of free Edans.

Inner-filter effect corrections were applied to the kinetics measurements according to Liu et al. [32]. As saturation could be achieved, kinetic constants (V_max_ and K_m_) were derived by fitting the corrected initial velocity to the Michaelis-Menten equation, *V* = *V*_max_ × [*S*]*/*(*K*_m_ + [*S*]), using the GraphPad Prism 7.0 software. k_cat_/K_m_ was calculated according to the equation *k*_cat_*/K*_m_ = *V*_max_*/*([*E*] × *K*_m_). Triplicate experiments were performed for each data point; the k_cat_, K_m_, and k_cat_/K_m_ values were presented as mean ± standard deviation (SD).

For the determination of the IC_50_, 50 nM of SARS-CoV-2 M^pro^ or O-M^pro^, or 100 nM P132H+T169S mutant was incubated with **13b-K** at various concentrations from 0 to 100 µM in reaction buffer at 37^◦^C for 10 min. Afterwards, the FRET substrate at a final concentration of 10 µM was added to each well, at a final total volume of 100 µL, to initiate the reaction. The GraphPad Prism 7.0 software (GraphPad) was used for the calculation of the IC_50_ values. Measurements of inhibitory activity of **13b-K** were performed in triplicate and are presented as the mean ± SD.

### Crystallization of the SARS-CoV-2 M^pro^ mutants

A freshly purified protein solution at 15 mg/mL concentration was centrifuged at 12,000 x *g* and used to crystallize M^pro^ mutants as a free enzyme and in complex with **13b-K**. For the co-crystallization of the **13b-K**/M^pro^ complex, the protein was mixed with **13b-K** (dissolved in 100% DMSO) at a molar ratio of 1:5. Then, the mixture was incubated at 4°C overnight. The next day, centrifugation was applied (12,000 x *g*) to remove any precipitate. Subsequently, four commercially available screening kits, PACT premier, SG1 (ShotGun), Morpheus, and Ligand-Friendly Screen (LFS) from Molecular Dimensions, were used for crystallization using a Crystal Phoenix robot (Art Robbins). The sitting-drop vapor-diffusion method was used at 25°C, and 0.3 μL of protein solution were mixed with 0.3 μL of the reservoir and left to equilibrate against a 40-μL reservoir solution. Crystals appeared within three days under several conditions. First, the free enzyme crystals of O-M^pro^ were obtained from 200 mM NaF, 20% PEG 3350, and 10% ethyleneglycol at pH 8.2. Next, **13b-K** co-crystals were obtained under two different conditions at two different pH values: (1), 25% PEG3350 and 100 mM Bis-Tris pH 6.5, and (2), 200 mM MgCl_2_, 25% PEG 3350, and100 mM Tris pH 8.5. The free-enzyme crystals of P132H+T169S M^pro^ were obtained from 100 mM SPG (Succinic acid, Sodium phosphate monobasic monohydrate, Glycine), 25% PEG 1500, pH 8.0. The **13b-K** co-crystals with P132H+T169S M^pro^ were also obtained under two different conditions: (1), 200 mM NaF, 20% PEG 3350, 100 mM Bis-Tris propane, pH 7.5, and (2), 100 mM CHES (N-cyclohexyl-2-aminoethanesulfonic acid), 20% PEG 8000, pH 9.0. Crystals were fished from the drops and cryo-protected by mother liquor plus varied concentrations of glycerol (10%-20%). Subsequently, fished crystals were flash-cooled in liquid nitrogen.

### Diffraction data collection, phase determination, model building, and refinement

All diffraction data sets were collected using synchrotron radiation of wavelength 1.033 Å at beamline P11 of DESY (Hamburg, Germany), using an Eiger 2X 16M detector (Dectris) [33]. Five data sets of the mutated SARS-CoV-2 M^pro^, the free enzymes of O-M^pro^ and P132H+T169S mutants, as well as three datasets in complex with **13b-K**, were collected from the crystals grown under the aforementioned conditions. XDSapp [34], Pointless [35, 36], and Scala [36] were used for processing the datasets.

The datasets of the free enzymes of O-M^pro^ and P132H+T169S mutants were processed at resolutions of 1.91 Å and 1.80 Å, respectively, in monoclinic space group *C2* with one protomer in the asymmetric unit. Datasets of the complexes with **13b-K** were processed at resolutions of 2.3 Å and 2.48 Å for the O-M^pro^ crystals grown at pH 8.5 and 6.5, respectively, in triclinic space group *P1* (four protomers in the asymmetric unit). A dataset for the **13b-K** complex of P132H+T169S mutant was collected at resolution of 1.80 Å in monoclinic space group *P2_1_* (two protomers in the asymmetric unit). Molrep [37] from the CCP4 suite [38] was used for molecular replacement using coordinate set 6Y2E [9] as a starting model. eLBOW [39] from the Phenix suite [40] was employed for the generation of the geometric restraints for **13b-K**, and the inhibitor was built into the F_o_-F_c_ density by using the Coot software [41]. The initial rounds of refinement of the five structures were performed with Refmac5 [42]. Then, the initial models were refined further using Phenix [40], after adding the solvent and the inhibitor, **13b-K**. Statistics of diffraction data processing and the model refinement are given in Table S1.

## Computational methods and protocols

### All-atom PME continuous constant-pH molecular dynamics (CpHMD) titration simulations

To determine the protonation states of O-M^pro^, we performed titration simulations using the newly implemented all-atom PME CpHMD [22] method in the Amber22 program [23] and the asynchronous pH-replica exchange protocol [24] to enhance sampling and accelerate convergence. Starting from the pH 8.5 X-ray structure of O-M^pro^ (with the inhibitor removed), the LEAP utility in Amber22 [23] was used to add hydrogens and dummy hydrogen for all Asp, Glu, and His sidechains and construct the free N– and C-termini. The protein was solvated in an octahedral water box with a minimum distance of 11 Å between the protein heavy atoms and water oxygen atoms at the edges of the box. Sodium and chloride ions were added to neutralize the system (with titratable sites in the default or model protonation states) and reach an ionic strength of 150 mM. The ff14SB force field [43] and TIP3P model [44] were used to represent the protein and water, respectively. The system underwent energy minimization with the steepest descent for the first 200 steps and conjugate gradient algorithm for the last 300 steps; a harmonic restraint with the force constant of 100 kcal·mol^−1^·Å^−2^ was applied to the protein heavy atoms. With the same harmonic restraint, the system was heated from 100 K to 300 K using a Langevin thermostat for 100 ps with a time step of 1 fs. After heating, CpHMD was turned on and a two-step equilibration was performed at pH 7, whereby the harmonic force constant was 100 kcal·mol^−1^·Å^−2^ in the first 250 ps and 10 kcal·mol^−1^·Å^−2^ in the last 250 ps. Next, the system was further equilibrated in four steps at 21 pH conditions of the replica-exchange runs (see below), whereby the restraint force constant was reduced from 10 to 1, 0.1, and 0 kcal·mol^−1^·Å^−2^, and each step lasted 500 ps. The production run utilized 21 pH replicas in the pH range of 2.5 to 8 with an interval of 0.25 pH units. An exchange between adjacent pH replicas was attempted every 2 ps (1000 MD steps) and the simulation lasted 45 ns per pH replica, which resulted in an aggregate sampling time of 945 ns. During the equilibration and production runs, the temperature was maintained at 300 K using the Langevin thermostat with a collision frequency of 1 ps^-1^. The isotropic Berendsen barostat was used to maintain a pressure of 1 bar with a relaxation time of 1 ps. The particle-mesh Ewald (PME) method with a real-space cutoff of 8 Å and grid space of 1 Å was used to treat long-range electrostatic interactions. The SHAKE algorithm was used to constrain the bonds involving hydrogen atoms to allow the integration step of 2 fs. Except for His132 and Glu240, the protonation states of all Asp, Glu, and His at pH 7 are in agreement with our previous simulations of the WT-M^pro^ [28].

### Fixed-charge molecular dynamics (MD) simulations

The fixed-charge MD simulations of O-M^pro^ were performed using the Amber20 program [45]. The solvated protein system was prepared as in the all-atom CpHMD simulation above. The protonation states were fixed as determined by the all-atom CpHMD for the pH 7 condition. Neutral histidines were set in the HIE state. In the simulations (two independent runs) starting from our pH 8.5 X-ray structure, His132 was set in the neutral state. In the simulations (four independent runs) starting from 7TOB, His132 was set in the neutral (two independent runs) or charged (two independent runs) state. The system was first energy minimized under the harmonic restraints of 100 kcal·mol^−1^·Å^−2^ placed on the protein heavy atoms for a total of 20000 steps using the steepest descent (first 10000 steps) followed by 10000 steps using the conjugate gradient algorithm. The system was gradually heated from 0 K to 300 K in 1 ns under the same harmonic restraints. Next, five stages of equilibration (each 10 ns) were conducted, whereby the harmonic restraints were gradually reduced from 10 to 5, 2, 1, and 0.1 kcal·mol^−1^·Å^−2^. All other settings were the same as in the CpHMD simulations. Two 500-ns production runs were performed based on our pH 8.5 X-ray structure of O-M^pro^. Four 1-μs production runs were performed based on the X-ray structure 7TOB, two of them with His132(0) and two of them with His132(+).

### Empirical calculations of protein stability change upon mutation using Rosetta

The changes in the stability of the free M^pro^ dimer and monomer upon the P132H mutation were estimated separately using the ddG_monomer application in the Rosetta software suite [27]. In this method, an ensemble of structures of the P132H mutant was generated from an input WT structure (PDB ID 7VH8 [46]; nirmatrelvir was removed). The change in folding free energy was calculated as the difference in the energy scores (using the REF2015 energy function [47]) between the WT and P132H M^pro^ structures. The high-resolution protocol was followed, which allowed for both backbone and side-chain adjustment. Rosetta’s standard side-chain optimization module first optimized the structure, then three sequential minimization calculations were performed where the Lennard-Jones potential was scaled by 0.1, 0.33, and 1.0, respectively. This was repeated 50 times for both the WT and mutant proteins, then the average score for each system was calculated. To prevent the backbone from deviating from the initial structure, distance restraints were placed on the alpha carbon atoms following the default protocol [27].

## Acknowledgements

We thank the staff at synchrotron beamline P11, DESY, Hamburg, Germany (Johanna Hakanpää et al.) for their support and Yuri Kusov, as well as Judith Röske for discussion. Financial support from the German Center for Infection Research (DZIF; project FF 01.905, to R.H.) and the National Institutes of Health (R35GM148261 to J. S.) is gratefully acknowledged. R.H. is also supported by the Government of Schleswig-Holstein through its Structure and Excellence Fund as well as by a close partnership between the Possehl Foundation (Lübeck) and the University of Lübeck.

