## Supplemental Materials for "Why is the Omicron main protease of SARS-CoV-2 less stable than its wild-type counterpart? A crystallographic, biophysical, and theoretical study of the free enzyme and its complex with inhibitor 13b-K"

**Supporting InformationTable S1**. Diffraction data and model refinement statistics

**Table S1**. Diffraction data and model refinement statistics

| | P132H_Free_ Enzyme | P132H_13bK_ pH 8.5 | P132H_13bK_ pH 6.5 | P132H+T169S_ Free_Enzyme | P132H+T169S_ 13b-K |
|---|---|---|---|---|---|
| Wavelength (Å) | 1.033 | 1.033 | 1.033 | 1.033 | 1.033 |
| Resolution range (Å) | 43.99-1.91 (1.98-1.91) | 40.9-2.3 (2.38-2.30) | 44.39-2.48 (2.57-2.48) | 33.98 - 1.80 (1.86-1.80) | 41.44-1.70 (1.76-1.70) |
| Space group | C 2 | P 1 | P 1 | C 2 | P 21 |
| a b c (Å)<br>α β γ ( º ) | 45.08 53.20 112.83<br>90.00 100.32 90.00 | 45.49 71.46 85.80<br>85.61 80.43 78.65 | 45.85 71.80 85.76<br>84.72 80.37 78.39 | 113.746 53.384 44.80<br>90 102.38 90 | 45.54 64.03 103.69<br>90.00 90.96 90.00 |
| Total reflections | 39598 (3839) | 90223 (9230) | 72463 (7364) | 97099 (9801) | 227360 (22894) |
| Unique reflections | 20086 (1965) | 41704 (4250) | 33675 (3397) | 24113 (2382) | 64655 (6414) |
| Multiplicity | 2.0 (2.0) | 2.2 (2.2) | 2.2 (2.2) | 4.0 (4.1) | 3.5 (3.6) |
| Completeness (%) | 95.53 (85.48) | 89.79 (91.20) | 90.01 (90.90) | 98.58 (98.72) | 98.37 (98.27) |
| Mean I/σ (I) | 9.74 (1.60) | 6.96 (1.94) | 9.21 (1.77) | 19.85 (1.39) | 10.92 (1.28) |
| Wilson B-factor | 32.15 | 28.67 | 50.05 | 38.17 | 22.15 |
| Rmerge | 0.0233 (0.4049) | 0.09499 (0.4351) | 0.05711 (0.5032) | 0.03003 (0.9106) | 0.07188 (0.9213) |
| Rmeas | 0.03295 (0.5726) | 0.1281 (0.5867) | 0.07732 (0.6808) | 0.03462 (1.048) | 0.08483 (1.082) |
| Rpim | 0.0233 (0.4049) | 0.08536 (0.3908) | 0.05174 (0.4556) | 0.01696 (0.5124) | 0.04457 (0.5629) |
| CC1/2 | 0.999 (0.93) | 0.993 (0.774) | 0.997 (0.76) | 1 (0.592) | 0.998 (0.635) |
| CC* | 1 (0.982) | 0.998 (0.934) | 0.999 (0.929) | 1 (0.862) | 1 (0.881) |
| Reflections used in refinement | 19634 (1743) | 41634 (4249) | 33628 (3396) | 24108 (2382) | 64640 (6408) |
| Reflections used for Rfree | 969 (76) | 2021 (212) | 1627 (157) | 1184 (114) | 3318 (327) |
| Rwork | 0.1946 (0.3945) | 0.1991 (0.2675) | 0.2112 (0.3463) | 0.2105 (0.3250) | 0.1943 (0.3263) |
| Rfree | 0.2366 (0.4681) | 0.2492 (0.3645) | 0.2488 (0.3528) | 0.2557 (0.3467) | 0.2230 (0.3648) |
| CC(work) | 0.970 (0.549) | 0.958 (0.832) | 0.958 (0.841) | 0.961 (0.629) | 0.965 (0.828) |
| CC(free) | 0.964 (0.367) | 0.900 (0.695) | 0.940 (0.813) | 0.942 (0.524) | 0.954 (0.745) |
| RMS(bonds) | 0.008 | 0.008 | 0.009 | 0.009 | 0.008 |
| RMS(angles) | 1.09 | 1.06 | 1.13 | 1.7 | 1.04 |
| Ramachandran favored (%) | 97.03 | 97.03 | 96.29 | 98.03 | 98.36 |
| Ramachandran allowed (%) | 2.64 | 2.89 | 3.55 | 1.64 | 1.48 |
| Ramachandran outliers (%) | 0.33 | 0.08 | 0.17 | 0.33 | 0.16 |
| Rotamer outliers (%) | 1.49 | 0.48 | 1.23 | 2.66 | 0.38 |
| Clash score | 3.34 | 4.68 | 4.54 | 2.99 | 2.75 |
| Average B-factor | 39.32 | 30.21 | 56.9 | 45.17 | 27.30 |

**Table S2**. Summary of continuous constant pH and fixed-charge molecular dynamics (MD) simulations of the free O-M$^{pro}$

| PDB ID | Simulation type | Length | H132/E240 conformation in the X-ray structure |
|---|---|---|---|
| Free O-M$^{pro}$ (this work) | PME-CpHMD | 45.1 ns x 21 | Stacking |
| Free O-M$^{pro}$ (this work) | Fixed charge; His132(0) | 500 ns x 2 | Stacking |
| 7TOB | Fixed charge; His132(+) | 1000 ns x 2 | Stacking |
| 7TOB | Fixed charge; His132(0) | 1000 ns x 2 | Stacking |

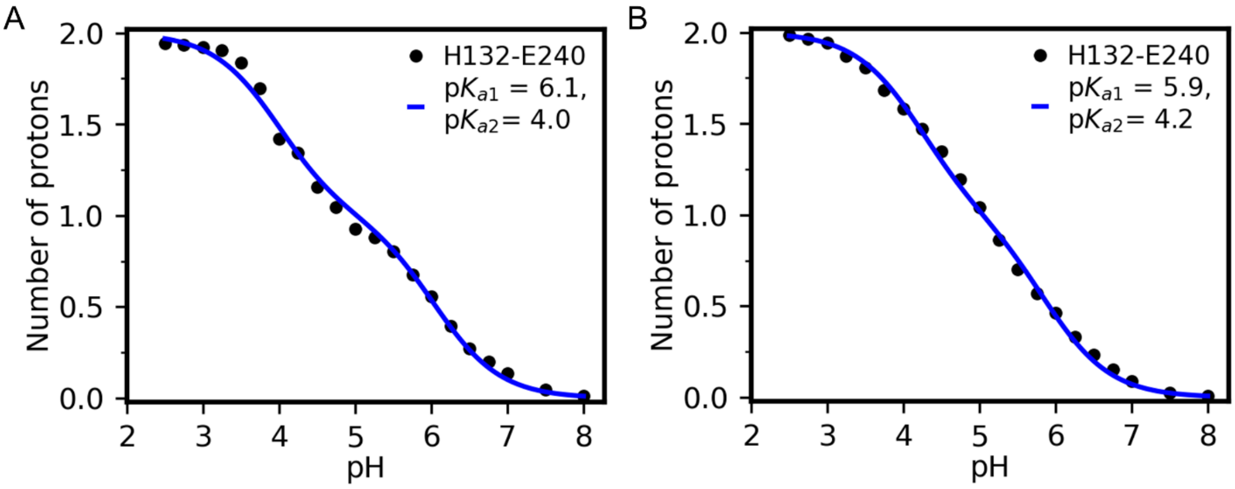


**Figure S1**. **Titration of His132 and Glu240 in the free O-M$^{pro}$ is coupled**. Average number of protons bound to the H132/E240 pair in the free O-M$^{pro}$ at different pH for protomer A (**A**) and B (**B**). The macroscopic stepwise p$K_a$ values were obtained by fitting to a coupled titration model [1, 2].

$$\langle P \rangle = \frac{10^{pK_{a2}-pH} + 2 \cdot 10^{pK_{a1}+pK_{a2}-2pH}}{1+10^{pK_{a2}-pH}+10^{pK_{a1}+pK_{a2}-2pH}} \quad \text{(S1)}$$

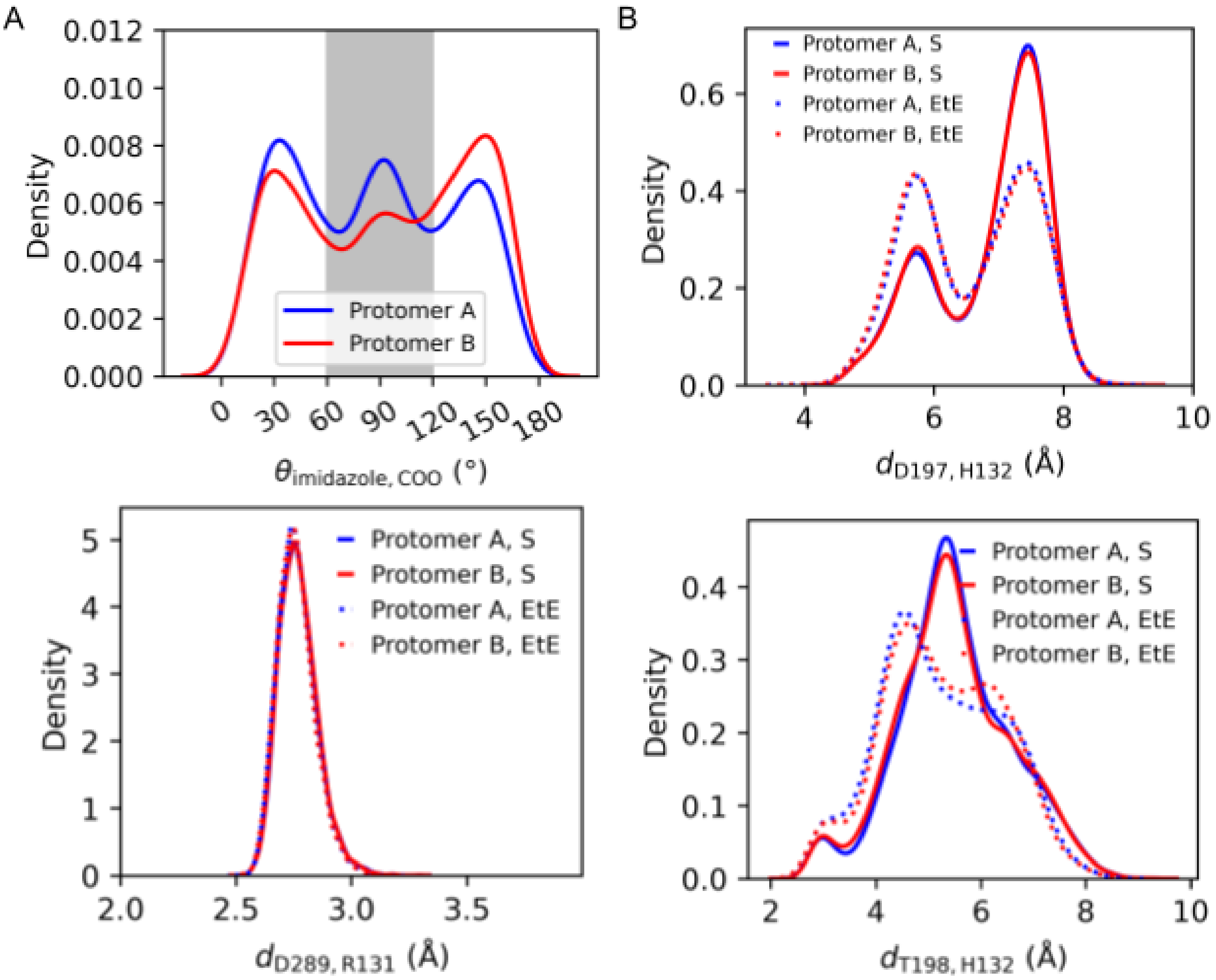


**Figure S2**. **The second fixed-charge simulation of the free O-M$^{pro}$ based on the free O-M$^{pro}$ structure of this work gave similar results. A**. distribution of the angle between the normal vectors of the H132 imidazole and E240 carboxylate planes for protomer A (blue) and B (red). The grey area represents the edge-to-edge (EtE) conformations while the other parts represent the stacking (S) conformations. **B**. Distribution of the minimum distance between the Asp197 carboxylate oxygens and the His132 backbone carbonyl oxygen. **C**. Distribution of the minimum distance between the Asp289 carboxylate oxygens and Arg131 guanidinium nitrogen atom. **D**. Distribution of the distance between the Thr198 hydroxyl oxygen and His132 imidazole ND atom. The data for the stacking (S, solid lines) and edge-to-edge (EtE, dotted lines) conformations of protomer A (blue) and B (red) are shown separately.

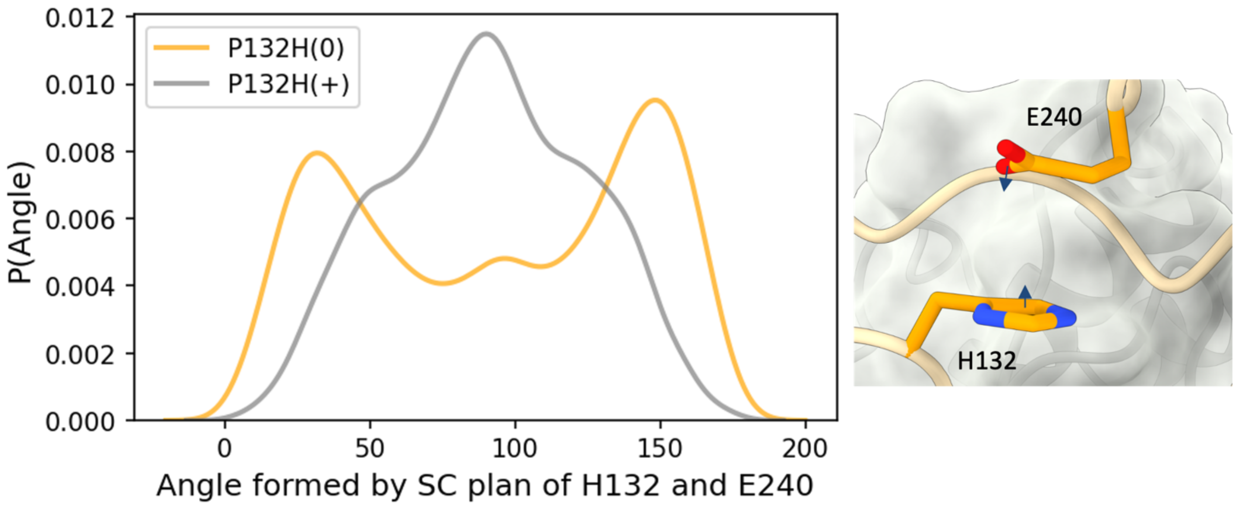


**Figure S3. Fixed-charge MD simulations based on the X-ray structure 7TOB (inhibitor removed) confirmed the preference of the stacking conformation by the neutral His132.** The distribution of the angle formed by the sidechain planes of His132 and Glu240 (same as the angle between the normal vectors). A zoomed-in view of His132 and Glu240 in the stacking conformation is given. The normal vectors of the two sidechain planes are shown.

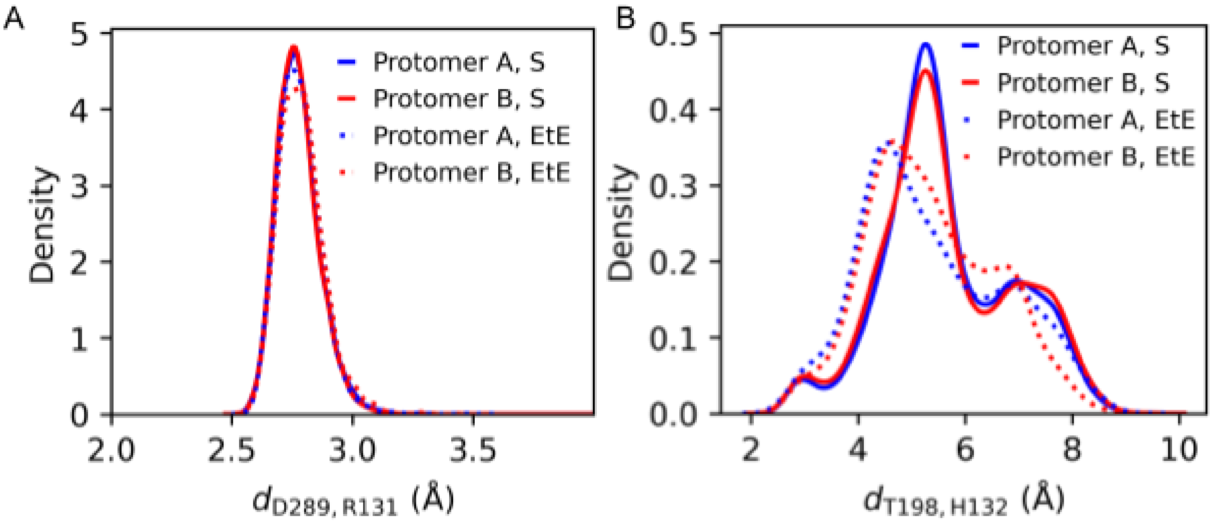


**Figure S4**. **Investigation of the Asp289-Arg131 salt-bridge and Thr198-His132 hydrogen bond interactions from the first run of the fixed-charge MD simulation of O-M$^{pro}$ based on the current X-ray structure of free O-M$^{pro}$**. **A**. Distribution of the minimum distance between the Asp289 carboxylate oxygens and Arg131 guanidinium nitrogens. **B**. Distribution of the distance between the Thr198 hydroxyl oxygen and the His132 imidazole ND atom. The data for the stacking (S) and edge-to-edge (EtE) conformations are represented as solid and dotted lines, respectively. Protomer A and B are shown in blue and red, respectively.

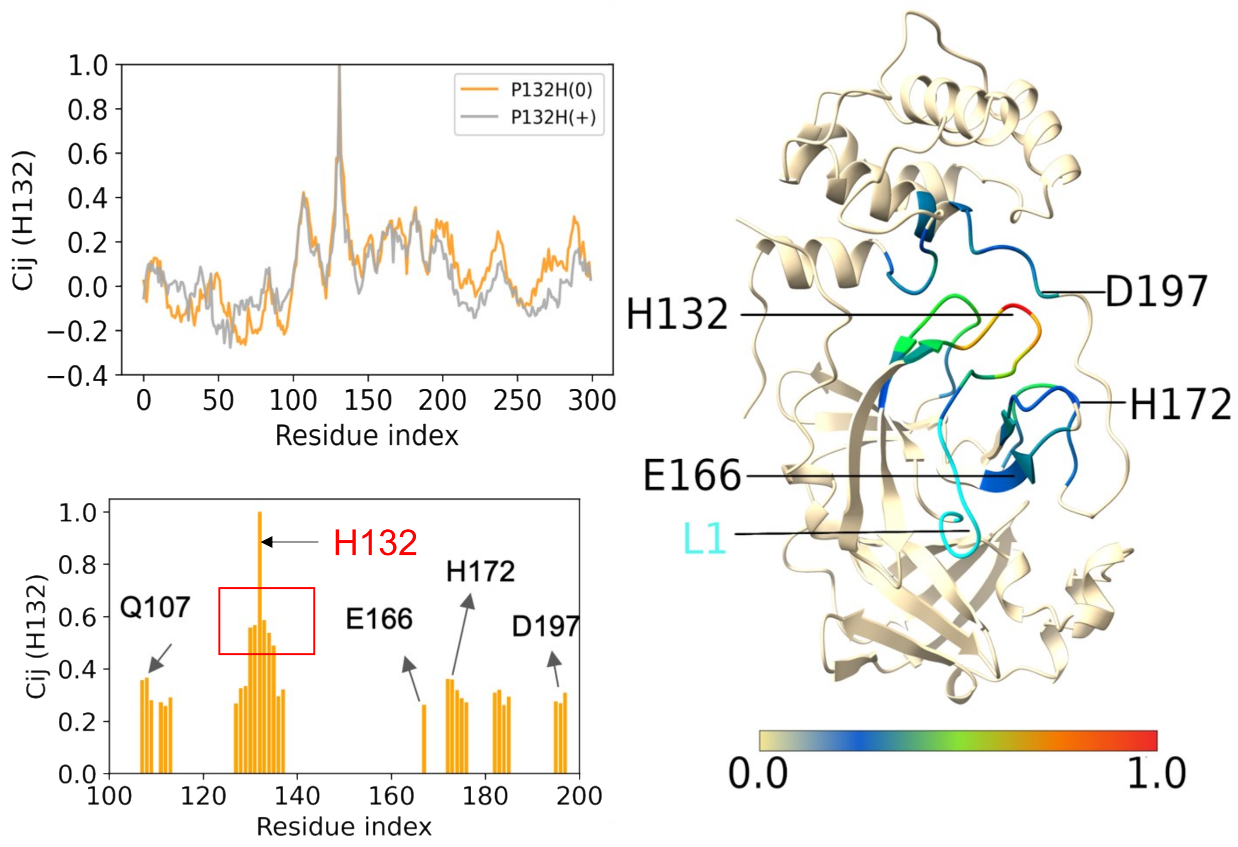


**Figure S5. Cross correlation analysis revealed dynamic correlation between His132 and residues in the domain II/III interface of the free O-M$^{pro}$. Left**. Cross correlation coefficients between His132 and all residues (Cα atoms) in the O-M$^{pro}$ dimer. The closed-up review of the correlation plot between residues 100 and 200 is given on the bottom. Only residues having coefficients above 0.25 are shown. A few residues are labeled to guide the eye to the sequence regions. The boxed region is where the cross-correlation coefficients exceed 0.5. **Right**. The correlation coefficients mapped onto the O-M$^{pro}$ monomer structure. The calculations used the last 500 ns of the fixed-charged MD simulations of the free O-M$^{pro}$ with the neutral His132(0) or charged His132(+). The values for the two protomers and two trajectories were averaged. The Bio3D package [3]. was applied.

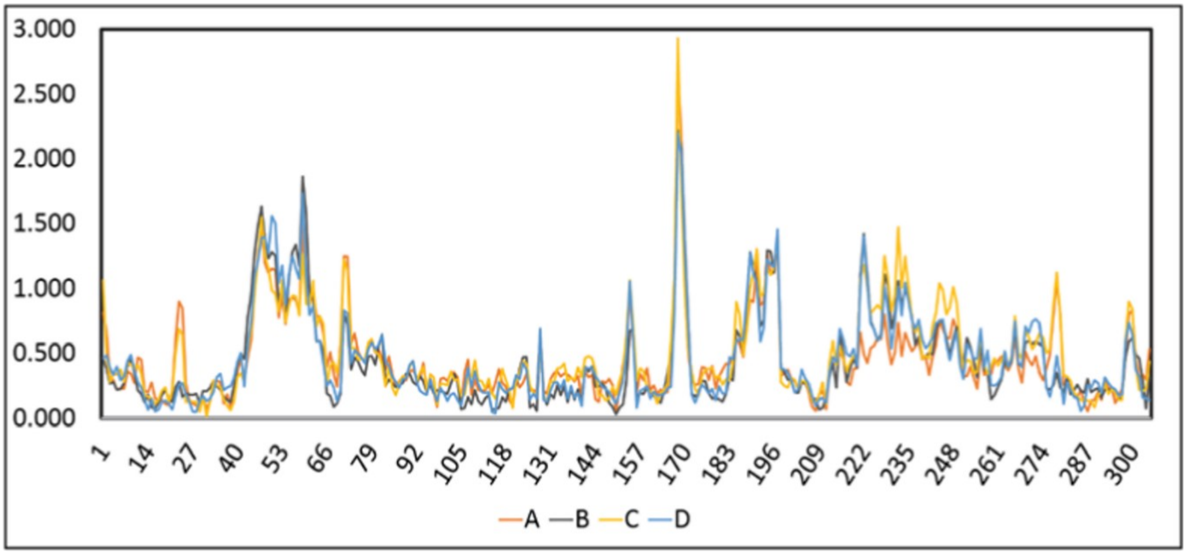


**Figure S6.** The RMSD values for the $C_\alpha$ of each amino acid of the four chains, A-D, of O-M$^{pro}$ model, depicted in orange, black, yellow, and blue, respectively. The values for panel C were computed from the crystallographic models using the built-in $rms_{cur}$ function in $PyMOL$.[4]

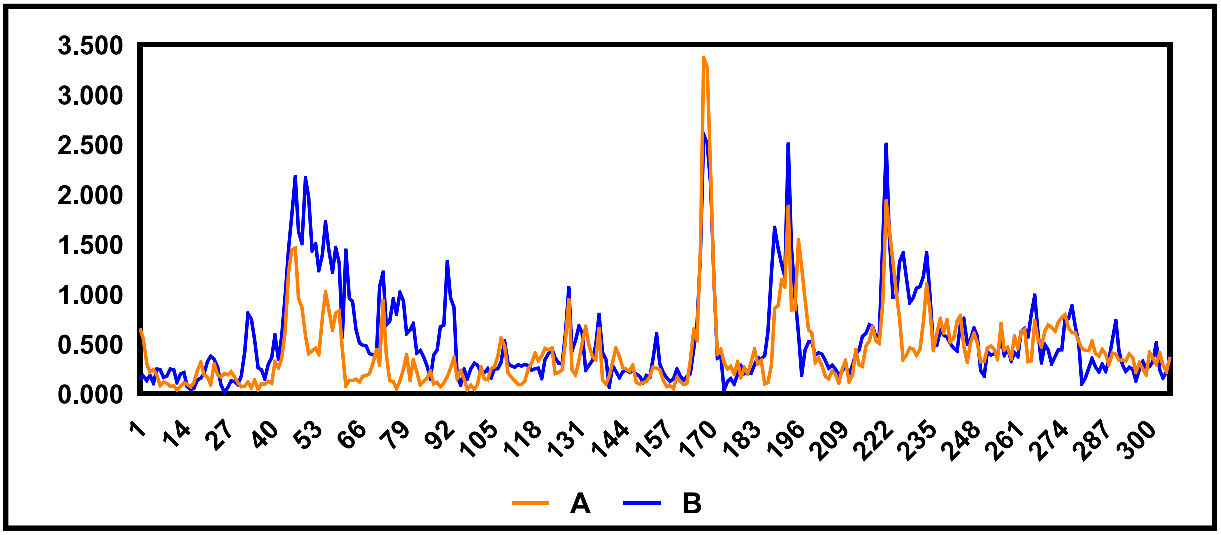


**Figure S7.** The RMSD values for the $C_\alpha$ of each amino acid of the two chains, A (orange) and B (blue), of M$^{pro}$-P132H+T169S model against the WT-M$^{pro}$ (6Y2E). The specific regions showing the high RMSD values are corresponding to the same regions in the O-M$^{pro}$ model.

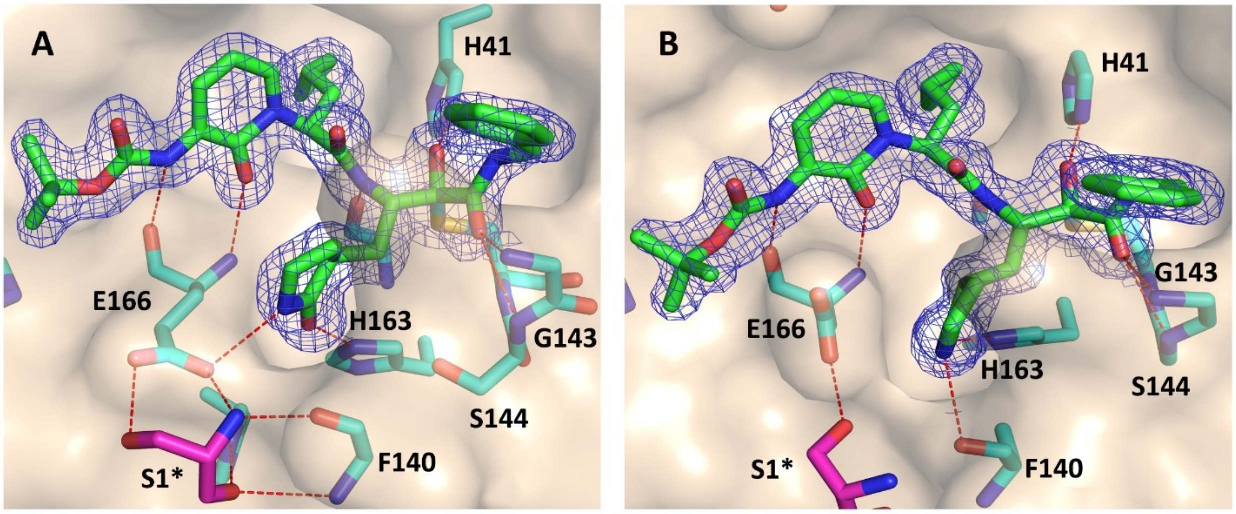


**Figure S8. X-ray structures of M$^{pro}$-P132H+T169S complexed with 13b-K under pH 9.0, in the substrate-binding cleft.** The 2F$_o$-F$_c$ map carved around the inhibitor is depicted as blue mesh at 1.0 σ. Polar contacts around inhibitor are represented as red dashed line. There are 2 copies found in this model, the Glu166 formed a salt bridge to Ser1 in promoter A (**A**) and formed a H-bond in promoter B (**B**), building a highly similar structure to M$^{pro}$-P132H+T169S structure at pH 7.5 (Fig. 5).
